# Hepatitis delta virus RNA decline post inoculation in human NTCP transgenic mice is biphasic

**DOI:** 10.1101/2023.02.17.528964

**Authors:** Stephanie Maya, Leeor Hershkovich, E Fabian Cardozo-Ojeda, Elham Shirvani-Dastgerdi, Jay Srinivas, Louis Shekhtman, Susan L Uprichard, Andrew R Berneshawi, Thomas R Cafiero, Harel Dahari, Alexander Ploss

## Abstract

**Background and Aims:** Chronic infection with hepatitis B and hepatitis delta viruses (HDV) is considered the most serious form of viral hepatitis due to more severe manifestations of and accelerated progression to liver fibrosis, cirrhosis, and hepatocellular carcinoma. There is no FDA-approved treatment for HDV and current interferon-alpha treatment is suboptimal. We characterized early HDV kinetics post inoculation and incorporated mathematical modeling to provide insights into host-HDV dynamics.

**Methods:** We analyzed HDV RNA serum viremia in 192 immunocompetent (C57BL/6) and immunodeficient (NRG) mice that did or did not transgenically express the HDV receptor - human sodium taurocholate co-transporting peptide (hNTCP).

**Results:** Kinetic analysis indicates an unanticipated biphasic decline consisting of a sharp first-phase and slower second-phase decline regardless of immunocompetence. HDV decline after re-inoculation again followed a biphasic decline; however, a steeper second-phase HDV decline was observed in NRG-hNTCP mice compared to NRG mice. HDV-entry inhibitor bulevirtide administration and HDV re-inoculation indicated that viral entry and receptor saturation are not major contributors to clearance, respectively. The biphasic kinetics can be mathematically modeled by assuming the existence of a non-specific binding compartment with a constant on/off-rate and the steeper second-phase decline by a loss of bound virus that cannot be returned as free virus to circulation. The model predicts that free HDV is cleared with a half-life of 18 minutes (standard error, SE: 2.4), binds to non-specific cells with a rate of 0.06 hour^-1^ (SE: 0.03), and returns as free virus with a rate of 0.23 hour^-1^ (SE: 0.03).

**Conclusions:** Understanding early HDV-host kinetics will inform pre-clinical therapeutic kinetic studies on how the efficacy of anti-HDV therapeutics can be affected by early kinetics of viral decline.

**LAY SUMMARY:** The persistence phase of HDV infection has been studied in some animal models, however, the early kinetics of HDV in vivo is incompletely understood. In this study, we characterize an unexpectedly HDV biphasic decline post inoculation in immunocompetent and immunodeficient mouse models and use mathematical modeling to provide insights into HDV-host dynamics. Understanding the kinetics of viral clearance in the blood can aid pre-clinical development and testing models for anti-HDV therapeutics.

## INTRODUCTION

Hepatitis delta virus (HDV) is a single-stranded RNA virus belonging to the *Kolmioviridae* family, genus Deltavirus that requires the hepatitis B virus (HBV) envelope proteins to package infectious virions and spread (*1, 2*). While it has been previously estimated that approximated 15 million are chronically infected with HDV a recent meta-analysis of epidemiological studies published between 1977 and 2016 suggests that this number may be closer to 60-70 million (*3*). This estimation revolves around the current understanding that HDV solely exploits HBV envelope proteins (*4-6*); however, recent data argues the possibility that HDV might utilize envelope proteins of a variety of viruses (*7-10*), although its implication in natural settings is still unknown (*11-13*). There is firm clinical data that suggests that co-infection with HDV and HBV results in significantly worse and accelerated progression to liver fibrosis, cirrhosis, and hepatocellular carcinoma than HBV infection alone (*14, 15*). Nevertheless, by utilizing HBV surface antigens (HBsAg) as its envelope proteins, HDV infects hepatocytes by first attaching to heparan sulfate proteoglycans (HSPGs) and subsequently binding to the HBV receptor, human sodium taurocholate co-transporting polypeptide (hNTCP) (*5, 16, 17*). Differences in the sequence between human NTCP and non-permissive species explain in part the limited host tropism of HBV and by extension HDV. While HDV, mediated through the preS1 region of the HBsAg can bind to mouse NTCP, differences in the amino acids 84-87 of the murine orthologue prevent HBV glycoprotein mediated uptake (*18*). Expression of human NTCP is sufficient to mediate HDV uptake and infection in mouse hepatocytes *in vitro (19)*, but these cells remain resistant to HBV (*20, 21*). These observations were corroborated in mice transgenically expressing hNTCP (*22, 23*) or a humanized allele of NTCP (*18*) that supports HDV. However, susceptibility was age-dependent and required inoculation with very high doses of HDV. We have subsequently shown that hNTCP transgenic (tg) mice support hepadnavirus glycoprotein-mediated uptake and persistent HDV infection when HBsAg is co-expressed (*23*).

The hNTCP tg mouse model allows us to investigate the kinetics of early HDV infection in mice. HDV long-term kinetics has been delineated in a few studies in humanized mice (*23-26*). Yet, HDV dynamics early during the infection, in particular the rate by which the virus enters hepatocytes or is cleared by the host, depending on host background immunity, remains to be defined.

In this study, we analyzed and mathematically modeled early HDV kinetics in hNTCP and non-hNTCP mice on an immunocompetent or immunodeficient background from inoculation until HDV viremia reached undetectable levels (lower limit of quantification (LLoQ)). To minimize the number of potentially confounding variables, we analyzed the kinetics of HDV infection in hNTCP mice in the absence of HBV or HBsAg. Although HDV requires a helper virus to propagate, evidence has shown that HDV can persist in human hepatocytes for at least 6 weeks (*27*) demonstrating that HBV is not needed for establishing intracellular HDV replication. Understanding early HDV dynamics such as the rate at which HDV can bind to hNTCP and be cleared from the bloodstream can create a baseline for pre-clinical therapeutic studies in these mouse models for teasing apart the effects of rapid HDV decline in the serum on anti-HDV treatments.

In the current study, we identified two distinct phases of viral decline, indicating a rapid initial virus decline followed by a slower decline phase until reaching LLoQ. Because of very frequent blood sampling, it was also possible to estimate the rate of HDV clearance from blood and explain the nature of the biphasic decline via mathematical modeling.

## MATERIALS AND METHODS

A detailed description of the Materials and Methods used in this study is included in the supplementary information.

## RESULTS

### HDV RNA in serum of hNTCP transgenic and non-transgenic mice undergoes a biphasic decline after single HDV inoculation

To characterize the early kinetics of HDV in mice, we first sought to delineate the stability of HDV in mouse serum after infection in the absence of HBV or ongoing production of HBV surface antigen. To this end, we utilized hNTCP transgenic or non-transgenic mice either on the C57BL/6 or NRG background, the latter lacking functional B, T and natural killer cells. These cohorts of mice were intravenously administered 1 × 10^9^ genomic equivalents (GE) of cell-culture produced HDV and bled every 2 hours for 24 hours (**Fig 1A**). Viral RNA was extracted from mouse serum and analyzed by RT-qPCR at each timepoint.

**Figure 1.**
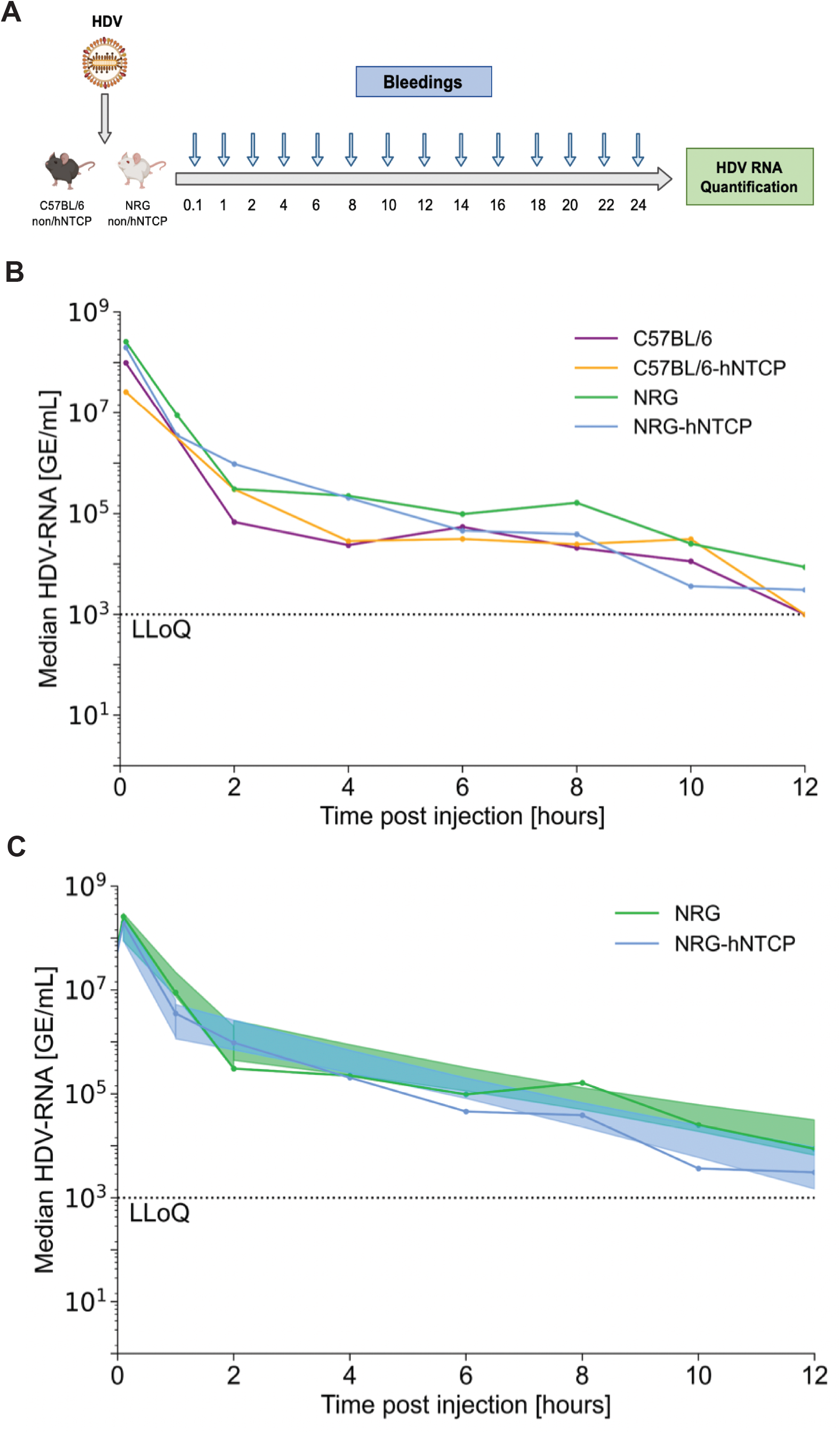
Single infection of HDV in C57BL/6 hNTCP or non-hNTCP mice and NRG hNTCP or non-hNTCP mice. (**A**) Schematic of C57BL/6 and NRG non-hNTCP or hNTCP mice infected with HDV and bled every two hours for the first 24 hours. Viral RNA was quantified from the serum by RT-qPCR. Schematic was created with Biorender.com. (**B**) Serum HDV RNA quantification over the first 14 hours of infection in C57BL/6 and NRG non-hNTCP or hNTCP mice. **(C)** Median serum HDV RNA for all HDV-naïve (first inoculation if applicable, including mice that were reinoculated at 4 and 12 hours, but their data is cut off before second inoculation) NRG and NRG-hNTCP mice, along with linear regressions and shaded 95% confidence intervals. LLoQ indicates the lower limit of quantification (1000 GE/mL). The second phase decline is significantly steeper in NRG-hNTCP mice than NRG mice (p=0.05, ANCOVA test). All data are represented as SEM. Each timepoint is representative of three mice.

HDV RNA levels in the singly-inoculated C57BL/6-hNTCP (n=18) and C57BL/6 (n=18) mice at 1 minute post infection (mpi) were not significantly different between the two experimental groups (**Fig 1B**). Thereafter, HDV RNA levels in the serum of C57BL/6-hNTCP mice followed a rapid decline within the first 4 hours post infection (hpi) followed by a slower decrease of serum HDV RNA which fell under the LLoQ by 14 hpi. In C57BL/6 mice, HDV viremia also declined swiftly in the first 4 hours followed by a decrease to LLoQ. Therefore, both C57BL/6 and C57BL/6-hNTCP mice followed a similar biphasic kinetic pattern characterized by a sharp first phase decline and a slower second phase clearance.

HDV RNA copy numbers in the inoculated NRG (n=30) and NRG-hNTCP (n=20) mice followed a similar biphasic decline within the first 4 hpi as their immunocompetent counterparts (**Fig 1B**). This rapid drop in HDV RNA levels was likewise followed by a slower decrease from 4 to 12 hpi, after which viremia plateaued at very low levels in both groups. Specifically, we observed that HDV RNA levels in NRG-hNTCP mice decreased more rapidly than in NRG mice in the first hour post infection, which was then followed by a slower decline (**Fig 1C)**. Overall, however, they both reached the LLOQ by 14 hpi.

Altogether, a biphasic viral decline was observed beginning with a sharp decrease in viral load 4 hpi in all mouse cohorts, followed by a slower second decline phase. Viremia in the immunocompetent mouse cohorts fell to the LLoQ by 8 hpi while viremia of the immunodeficient mice reached the limit of detection by 14 hpi.

### Viral entry inhibition by bulevirtide treatment in NRG-hNTCP mice had negligible effect on viral clearance

Since hepatocytes of NRG-hNTCP mice are permissive to HDV (*23*), we sought to determine whether the decline of virus in the blood circulation is due to viral binding to hNTCP. We employed treatment with bulevirtide (also known as Myrcludex B or Hepcludex), an HBV/HDV entry inhibitor (*23, 30, 31*), to block viral binding to hNTCP by competitive inhibition. NRG-hNTCP mice (n=9) were treated with bulevirtide 1 hour prior to inoculation with 1 × 10^8^ GE of HDV (**Fig 2A**). Untreated NRG (n=6) and NRG-hNTCP (n=6) mice were also injected with 1 × 10^8^ GE HDV per mouse and were bled immediately after inoculation followed by bleedings every 2 hours for 24 hours hpi. Viral HDV RNA was extracted from mouse serum and analyzed by RT-qPCR at each timepoint.

**Figure 2.**
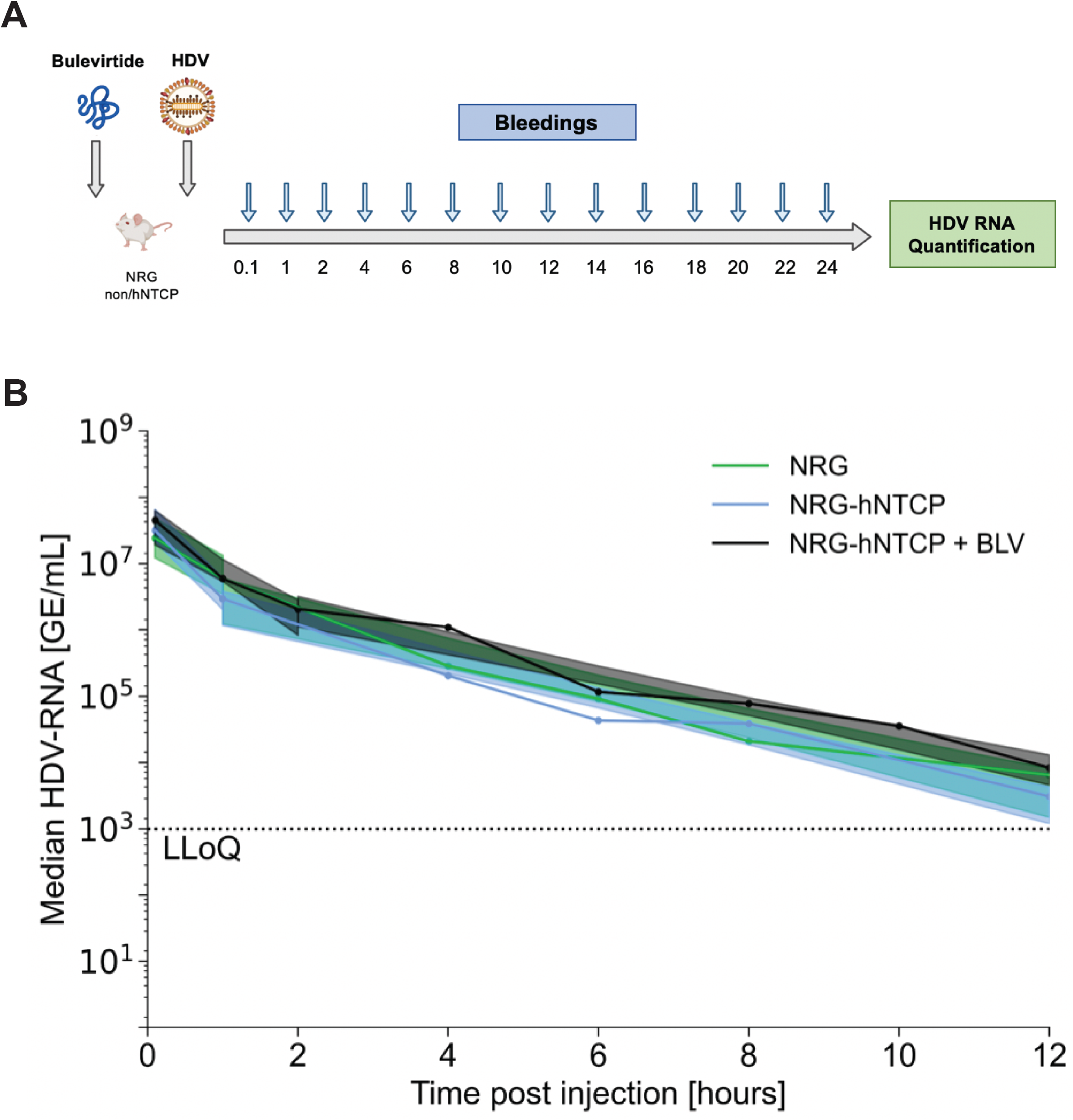
Bulevirtide treatment 1 hour prior to single infection of HDV in NRG hNTCP or non-hNTCP mice. (**A**) Schematic of NRG non-hNTCP or hNTCP mice pre-treated with bulevirtide and subsequently infected with HDV, followed by bleedings every two hours for the first 24 hours. Viral RNA was quantified from the serum by RT-qPCR. Schematic was created with Biorender.com. (**B**) Serum HDV RNA quantification over the first 12 hours of infection in and HDV-infected NRG, NRG-hNTCP, or bulevirtide-treated NRG-hNTCP (NRG-hNTCP + BLV) mice. Linear regression confidences are displayed as shaded 95% confidence intervals. LLoQ indicates the lower limit of quantification (1000 GE/mL). All data are represented as SEM. Each timepoint is representative of three mice.

In NRG-hNTCP mice treated with bulevirtide (bulevirtide-hNTCP), HDV RNA levels at 1 mpi were not significantly different between NRG-hNTCP and bulevirtide-hNTCP groups (**Fig 2B**). Thereafter, serum HDV levels in NRG-hNTCP mice followed a biphasic decline characterized by a rapid drop from 0 to 2 hpi (1.1 log/hr [95% CI: 0.77 -1.5]) and a slower decrease from 2 to 12 hpi 0.27 log/hr [95% CI: 0.23 -0.31]). The biphasic decline kinetics in bulevirtide-hNTCP mice similarly consisted of a rapid phase decline in the first 2 hours 1.0 log/hr [95% CI: 0.74 -1.3 and a slower phase until 12 hpi 0.25 log/hr [95% CI: 0.22 -0.28](**Fig 2B**). HDV RNA levels in NRG-hNTCP mice pre-treated with bulevirtide did display a small but detectable delay in clearance in the serum compared to untreated NRG-hNTCP mice (**Suppl Fig 2**). This could potentially be a result of a block in HDV uptake into hepatocytes due to bulevirtide competition; however, this observation is not statistically significant. The similar kinetics in mice on an NRG-hNTCP background, with or without bulevirtide, suggest the negligible effect of HDV binding on the decrease of virus in the bloodstream.

### Re-inoculation of HDV at 4 hpi in NRG hNTCP transgenic and non-transgenic mice results in similar biphasic declines

To further investigate whether receptor saturation or other binding site is the cause of this biphasic decline, we injected mice with HDV a second time at 4 or 12 hours following the initial inoculation, which characterized the end of each individual kinetic phase in single inoculation experiments. To observe the effect of re-inoculation at the end of the first phase, we utilized NRG (n=18) and NRG-hNTCP (n=15) mice and injected them with HDV at 1 × 10^8^ GE per mouse followed by re-inoculation at 4 hpi with the same viral load (**Fig 3A**).

**Figure 3.**
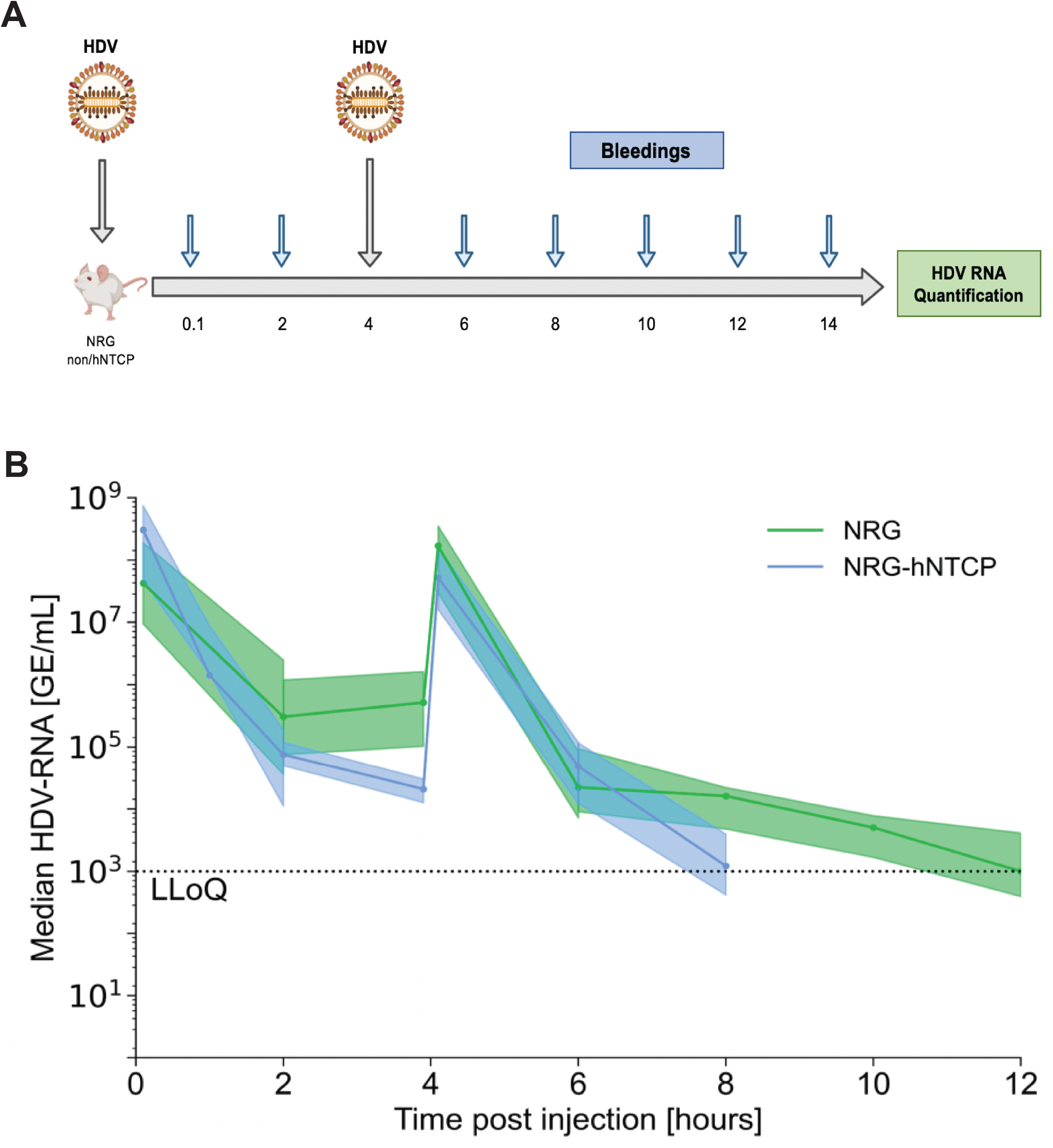
Re-infection of HDV in NRG-hNTCP mice 4 hours post initial infection. (**A**) Schematic of NRG and NRG-hNTCP mice infected with HDV at time zero and 4 hpi and bled every two hours for the first 24 hours. Viral RNA was quantified from the serum by RT-qPCR. Schematic was created with Biorender.com. (**B**) Median serum HDV RNA quantification over the first 12 hours of infection in NRG and NRG-hNTCP mice, along with linear regressions and shaded 95% confidence intervals. LLoQ indicates the lower limit of quantification (1000 GE/mL). All data are represented as SEM. Each timepoint is representative of three mice.

In doubly injected NRG and NRG-hNTCP mice, the HDV RNA levels followed a biphasic decline in accordance with a rapid drop from 0 to 2 hpi and a slower decrease from 2 to 4 hpi (**Fig 3B**). After re-infection, HDV RNA levels reached a peak around 1 × 10^8^ GE/mL which did not differ significantly from the initial RNA levels prior to re-infection. Afterwards, RNA levels followed a biphasic decline consistent with an initial rapid drop from initial levels in NRG mice and NRG-hNTCP mice. Notably, while the early rapid phase lasted 2 hours (from 4 to 6 hpi) post re-injection in NRG mice, the rapid phase in NRG-hNTCP mice lasted 4 hours post re-infection, from 4 to 8 hpi (**Fig 3B**). The rapid decline was then followed by the slower phase decrease in both mouse cohorts. Strikingly, levels of HDV RNA in the NRG-hNTCP mice decreased more rapidly after both HDV injections as compared to NRG mice. The second injection even resulted in clearance of viral RNA in the serum of NRG-hNTCP mice by 8 hpi while NRG mice experienced viral RNA clearance by 12 hpi.

### Re-inoculation of HDV at 12 hpi in NRG hNTCP transgenic and non-transgenic mice results slower second phase declines compared to 4 hpi re-inoculation

To probe the potential for receptor and/or other binging site saturation during the second phase decline of HDV, we re-inoculated the mice at 12 hpi. Thereby, we reasoned that we can determine whether there is a sharper or steadier decline in viral RNA when HDV is re-introduced during the second phase of decline. NRG (n=12) and NRG-hNTCP (n=15) mice were thus injected at 0 hpi and 12 hpi with bleedings every 2 hours for the first 24 hours (**Fig 4A**).

**Figure 4.**
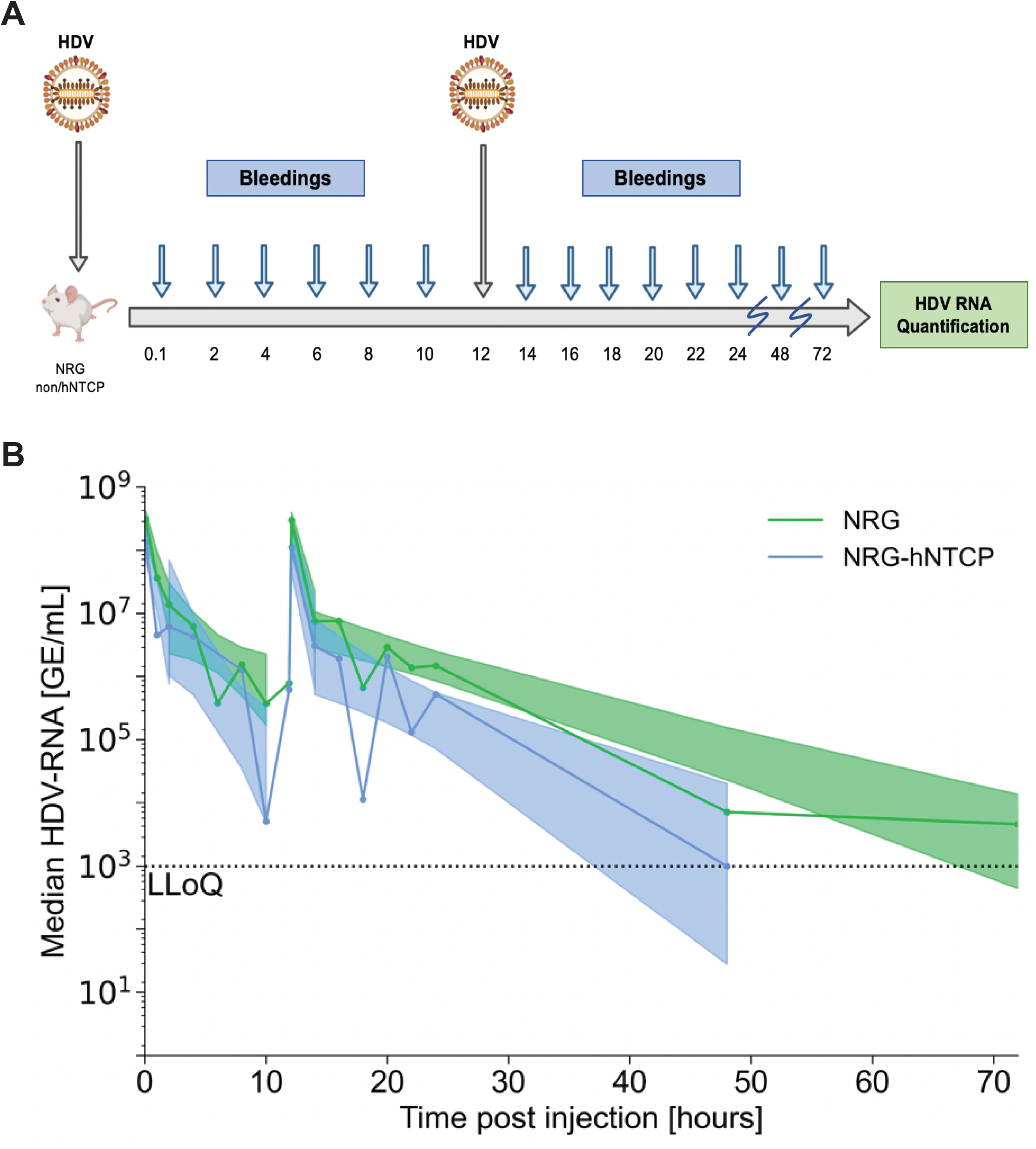
Re-infection of HDV in NRG and NRG-hNTCP mice 12 hours post initial infection. (**A**) Schematic of NRG and NRG-hNTCP mice infected with HDV at time zero and 12 hpi and bled every two hours for the first 24 hours. Viral RNA was quantified from the serum by RT-qPCR. Schematic was created with Biorender.com. (**B**) Serum HDV RNA quantification over all 72 hours of infection in NRG and NRG-hNTCP mice and linear regressions with shaded 95% confidence intervals. LLoQ indicates the lower limit of quantification (1000 GE/mL). The second phase decline after the second-inoculation is significantly steeper in NRG-hNTCP mice than NRG mice (p=0.02, ANCOVA test). Slope comparisons for the first inoculation are presented in Figure (1). All data are represented as SEM. Each timepoint is representative of three mice.

In both cohorts, HDV RNA biphasic decline comprised of a rapid drop from 0 to 4 hpi and a slower phase decline from 4 to 12 hpi (**Fig 4B**). The biphasic decline kinetics in both groups consisted of a rapid phase decline and slower phase decline until 12 hpi, after which the mice were re-infected with a second HDV inoculation dose. After re-injection, median HDV RNA levels reached a peak that was not significantly different from the RNA levels at the initial injections. Subsequently, HDV RNA levels followed a biphasic decline composed of an initial drop of RNA levels followed by the slower phase decrease. Remarkably, the HDV levels in the NRG mice did not fall below the LLoQ by 48 hpi as in the NRG hNTCP mice; instead, the levels remained elevated until 72 hpi (**Fig 4B**). Similar to the 4 hpi injections in **Fig 3B**, the NRG-hNTCP mice resulted in faster declines of HDV RNA in the serum following both injections compared to NRG mice. To analyze the potential impact of early immune responses, we also conducted the 12 hpi re-inoculation in transgenic and non-transgenic C57BL/6 mice. Comparatively, RNA levels in C57BL/6 and C57BL/6-hNTCP mice followed similar kinetics to NRG and NRG-hNTCP mice (**Suppl. Fig 1A and B**).

Overall, a biphasic decline before/after re-inoculation was observed in both 4 hpi and 12 hpi cohorts for transgenic and non-transgenic NRG mice (**Figs 3B and 4B**). Particularly, HDV RNA levels in NRG-hNTCP mice declined more rapidly following double HDV injections as compared to NRG mice. These data suggest that the hNTCP receptors for HDV binding are not saturated during early infection.

### Agglomerate kinetics analysis reveals faster clearance NRG-hNTCP mice compared to NRG mice after re-inoculations

To assess the overall effects and trends of the hNTCP receptor, immunocompetence, and of bulevirtide on HDV viral kinetics, mice were agglomerated for further analysis across experiment runs and cohorts.

Upon first intravenous injection of HDV in HDV-naïve mice (data from re-inoculated mice was cut off at the time of re-inoculation), all mouse strains followed a similar biphasic decline (**Fig 1B**). HDV RNA rose to a median 8.3 log GE/mL (interquartile, IQR 7.6-8.6). NRG mice experienced a rapid initial median decline in HDV RNA of 1.2 log/hr until 2 hpi, followed by a slower median decline of 0.19 log/hr [95% CI: 0.13-0.25] until 14 hpi. NRG-hNTCP mice experienced a similar rapid initial median decline in HDV RNA of 1.9 log/hr until 1 hpi, followed by a slower second median decline of 0.26 [95% CI: 0.20-0.31] log/hr (**Fig 1C and Table 1**).

**Table 1.**
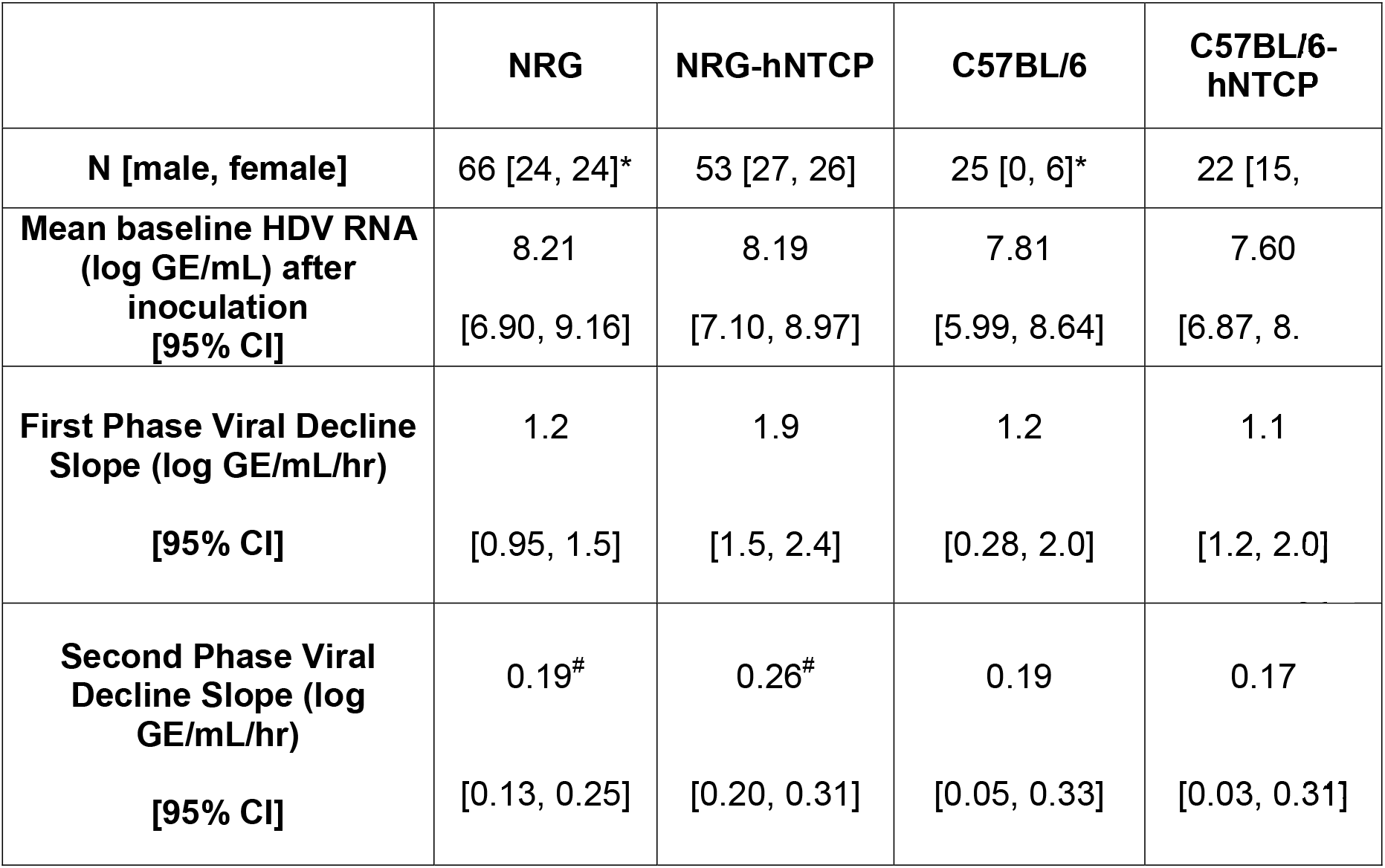
Summary of first inoculation agglomeratemo use kinetics for all mice, excluding mice that were administered bulevirtide. Data represented includes all HDV-naïve mice, including those that only received 1 inoculation, and those that were re-inoculated (including at 4 and 12 hours), but their data is cut-off before the second inoculation. Each mouse strain studied is represented as a single agglomerate distribution of all re-inoculation-time groups. The 95% confidence intervals (CI) for each decline are shown, demonstrating the steeper second-phase decline in NRG-hNTCP mice. *: Numbers in brackets represent known sex of mice within this subpopulation, the remainder are unspecified. ^#^: These slopes are significantly different from each other (p=0.05).

The HDV RNA kinetics of C57BL/6 and C57BL/6-hNTCP mice did not differ significantly between each other, or between their immunocompromised counterparts (NRG/NRG-hNTCP) (**Fig 1B**). Similarly, administration of bulevirtide to NRG-hNTCP mice did not affect the slope of decline compared to untreated NRG and NRG-hNTCP mice among the same cohort (**Fig 2B and Table 1**). Due to their negligible effects, and their smaller sample sizes, C57BL/6 and C57BL/6-hNTCP mice were precluded from mathematical modeling.

Following the re-inoculations at 4 or 12 hpi of NRG and NRG-hNTCP, the median HDV RNA rose to 8.14 log GE/mL (IQR: 7.78-8.59). Just as in the first inoculation, all mice showed a biphasic HDV RNA decline post-reinoculation. For both NRG and NRG-hNTCP mice, the viral kinetics of mice reinoculated at 4 hpi differed from those reinoculated at 12 hpi (**Fig 3B and 4B**). The NRG mice reinoculated at 4 hpi experienced a median rapid first-phase decline in HDV RNA of 1.9 log/hr [95% CI: 1.5-2.3] for 1 hour, followed by a slower decline of 0.23 [95% CI: 0.09-0.36] log/hr to LLoQ. The NRG mice that were reinoculated at 12 hpi showed a comparatively slower rapid first phase of HDV RNA/ decline of 0.79 [95% CI: 0.5-1.1] log/hr for 3 hours, followed by a slower second phase decline of 0.06 [95% CI: 0.04-0.07] log/hr. The NRG-hNTCP mice that were reinoculated at 4 hpi experienced a median rapid first-phase HDV RNA decline of 1.6 [95% CI: 1.3-2.0] log/hr for 1 hour, followed by a slower decline of 0.73 [95% CI: 0.39-1.1] log/hr. The NRG-hNTCP mice that were reinoculated at 12 hpi had a median rapid first-phase HDV RNA decline of 0.82 [95% CI: 0.39-1.2] log/hr for 3 hours, followed by a slower second phase of 0.10 [95% CI: 0.05-0.15] log/hr (**Table 2**).

**Table 2.**
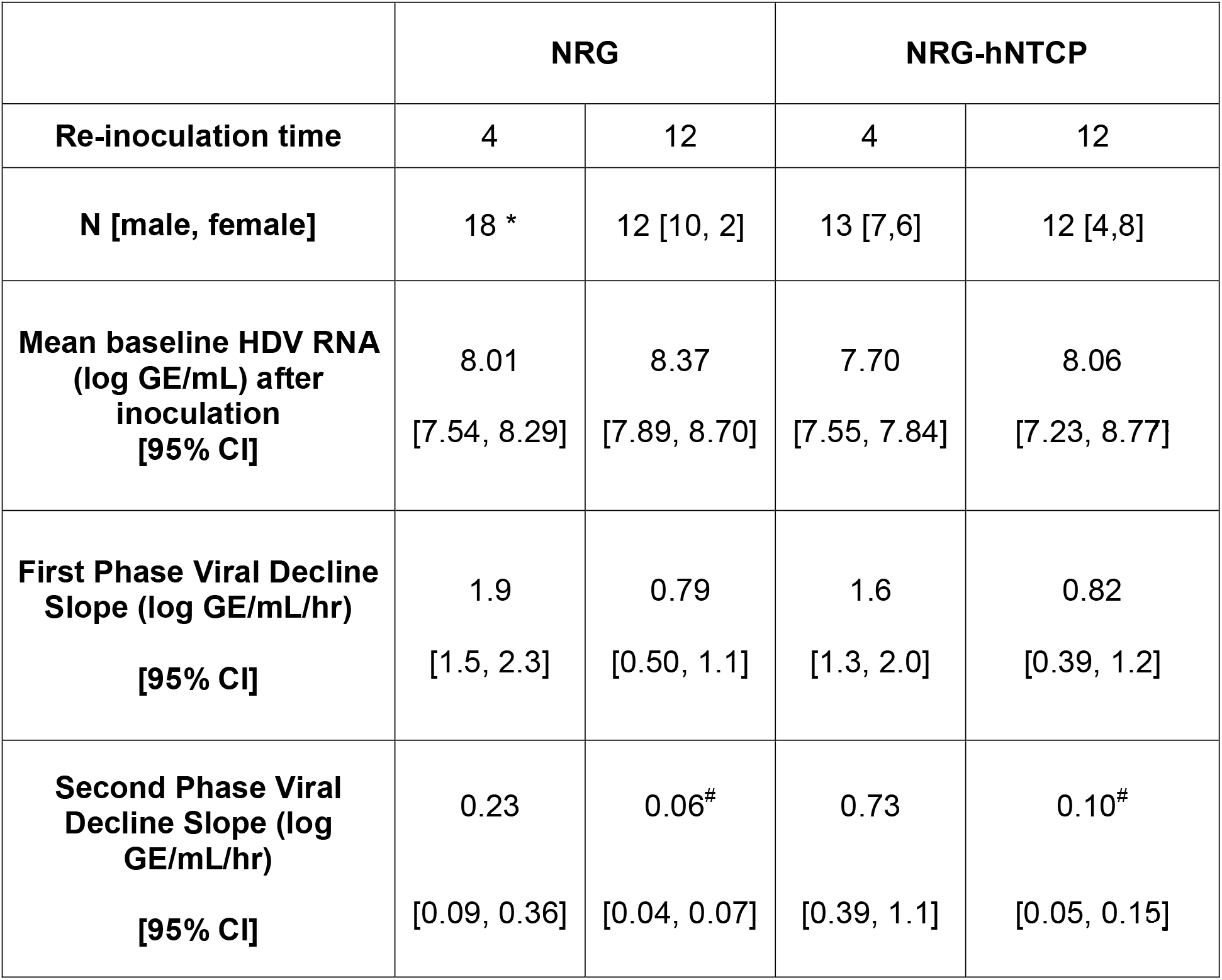
Summary of NRG and NRG-hNTCP second inoculation agglomerate kinetics. For both NRG and NRG-hNTCP, the 4-hour re-inoculation and 12-hour re-inoculation groups are represented separately. 95% confidence intervals for each decline are shown, demonstrating the steeper second-phase decline in NRG-hNTCP mice.*: Numbers in brackets represent known sex of mice within this subpopulation, the remainder are unspecified. ^#^: Significantly different slopes from each other (p=0.002).

A steeper second-phase was observed in NRG-hNTCP mice relative to NRG, both after the first inoculation (p=0.05, **Fig 1C**) and after the second inoculation (p=0.002, **Fig 4B**). The consistent steeper second-phase suggests that there may be a heightened immune response or receptor-bound virus in the second-phase decline for NRG-hNTCP mice. To provide insights into HDV-host kinetics in the NRG and NRG hNTCP mice, we developed a mathematical model as described next.

### A binding compartment mathematical model can be used to explain the biphasic viral decline in mice

Assuming no viral production and negligible effect of HDV entry on viral decline, a binding-compartment model (**Fig 5A and 5B**) was built to describe the experimental observations from the NRG and NRG-hNTCP mice. The model considers the dynamics of two populations of virus, the free-roaming virions, *v*_*f*_, and cell-bound HDV, *v*_*b*_. Assuming no production of new virions, free virus enters circulation by being released from a bound cell with rate-constant *k*_*on*_ and is removed from circulation at a general clearance rate *c*, as well as by being bound to a cell at rate-constant *k*_*on*_. The cell-bound *v*_*b*_ population grows as *v*_*f*_ is bound to a cell with rate-constant *k*_*on*_ and shrinks as *v*_*b*_ is released from the binding cell at rate-constant *k*_*off*_ or as it is lost (with no unbinding) with rate *k*_ℓ_ only in the transgenic NRG-hNTCP mice (**Fig 5A**). To account for the infection or re-infection of HDV at times 0, 4 or 12 hours, the function *R*(*t*) was incorporated (**Fig 5B**). Here, *R*(*t*) is a pulse function with form 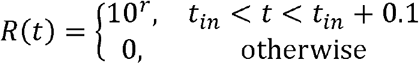, with *t*_*in*_ the time of inoculation or re-inoculation of HDV (0, 4 or/and 12 hours) and *r* is the orders of magnitude by which cell free HDV increases during inoculation or re-inoculation of virus.

**Figure 5.**
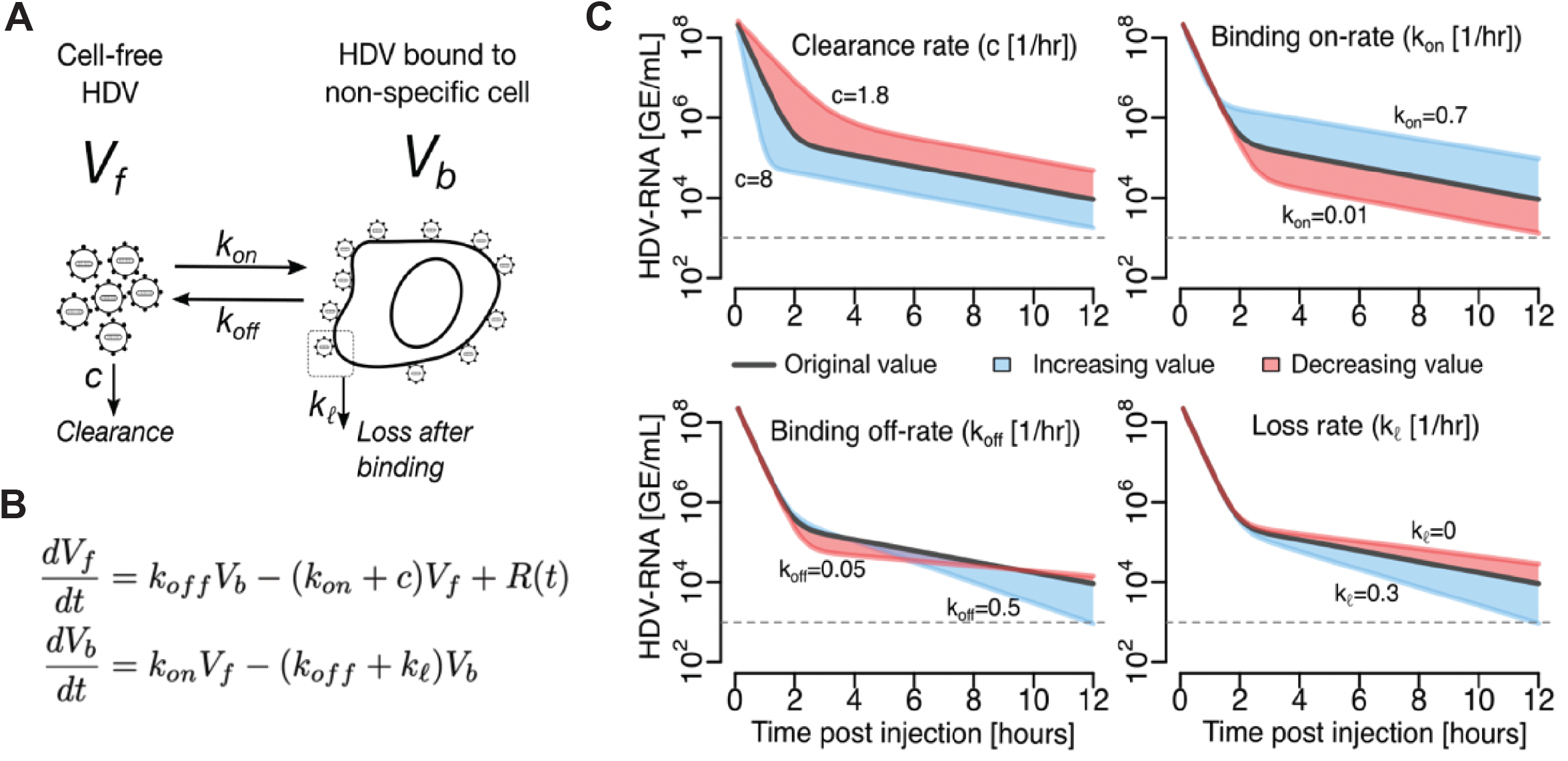
Binding compartment mathematical modeling. (**A**) Schematic of mathematical model: *v*_*f*_ is the free-roaming virions and *v*_*b*_ the cell-bound HDV. Free virus enters circulation by being released from a non-specific binding cell with rate-constant *k*_*on*_ and is removed from circulation at a general clearance rate *c*, as well as by being bound to a cell at rate-constant *k*_*on*_. The cell-bound *v*_*b*_ shrinks as it is released from the binding cell at rate-constant *k*_*off*_ or as it is lost with rate *k*_ℓ_ only in the transgenic NRG-hNTCP mice. (**B**) Model equation. Here, *R*(*t*) is a pulse function with form 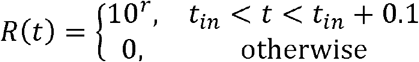 with *t*_*ln*_ the time of inoculation or re-inoculation of HDV (0, 4 or/and 12 hours) and *r* is the orders of magnitude by which cell free HDV increases during inoculation or re-inoculation of virus. (**C**) Model simulations indicate that the speed and time of the first phase of decline are primarily affected by the HDV clearance rate (top-left) and the binding on-rate (top-right). The second phase of decline is primarily affected by the binding off-rate (bottom-left) and internalization rate parameters (bottom-right). Model parameters in each graph are fixed at: *k*_*on*_ = 0.07 day^-1^, *k*_*off*_ = _ℓ_ 0.22 day^-1^, *k*_ℓ_ = 0.1 day^-1^, *c* = 3.71 day^-1^ (black curve). Adjustments made to each parameter are shown in red (underestimate) and blue (overestimate), for illustrative purposes.

To examine the model (**Fig 5A and 5B**) sensitivity to model parameters, a one-way sensitivity analysis was conducted (**Fig 5C**). The cell-free virus (*v*_*f*_) clearance rate constant (*c*) was positively associated with the slope and duration of the first phase of *v*_*f*_ decline, and the binding rate constant (*k*_*on*_) was inversely correlated with the duration of the first phase of HDV decline. The second phase of *v*_*f*_ decline is primarily associated with the loss and off-rate constant after binding (i.e., a larger binding compartment internalization and off-rate is associated with a faster second phase decline rate). The loss and off-rate constant (*k*_ℓ_ and *k*_*off*_, respectively) were positively associated with the slope of the second phase decline. Similarly, simulations predict that the majority of *v*_*f*_ rapidly become cell-bound (*v*_*b*_), which reach peak values followed by *v*_*b*_ decline (**Suppl. Fig 2**). The peak in *v*_*b*_ is inversely correlated with changes in clearance rate (*c*) and positively correlated with the binding rate constant (*k*_*on*_). The second phase decline rate was governed by the loss and off-rate constant (*k*_ℓ_ and *k*_*off*_, respectively), where a larger binding compartment loss and off-rate is associated with a faster second phase decline.

### Modeling suggests NRG-hNTCP mice experience a loss of bound HDV that does not return to circulation as free virus

We simultaneously fit this model to the NRG and NRG-hNTCP mouse data using a nonlinear mixed effect approach. We compared a model where *k*_ℓ_ *> 0*, i.e., bound virus in NRG-hNTCP mice gets lost leading to a faster second phase with respect to a null model where *k*_ℓ_ *= 0*. Using model selection, we found that a model assuming *k*_ℓ_ *>0* is more parsimonious to explain the data (ΔAICc=7.3). Examples of the model fits to viral load from NRG and NRG-hNTCP mice are shown in **Fig 6**. Model fits to HDV concentration to all mice are shown in **Supp. Fig 3, 4** using the individual parameters estimates in **Supp. Tables 1, 2** and the maximum likelihood estimates of the population distributions are shown in **Table 3** (for more details on the model parameter definitions see **Methods** section).

**Table 3.**
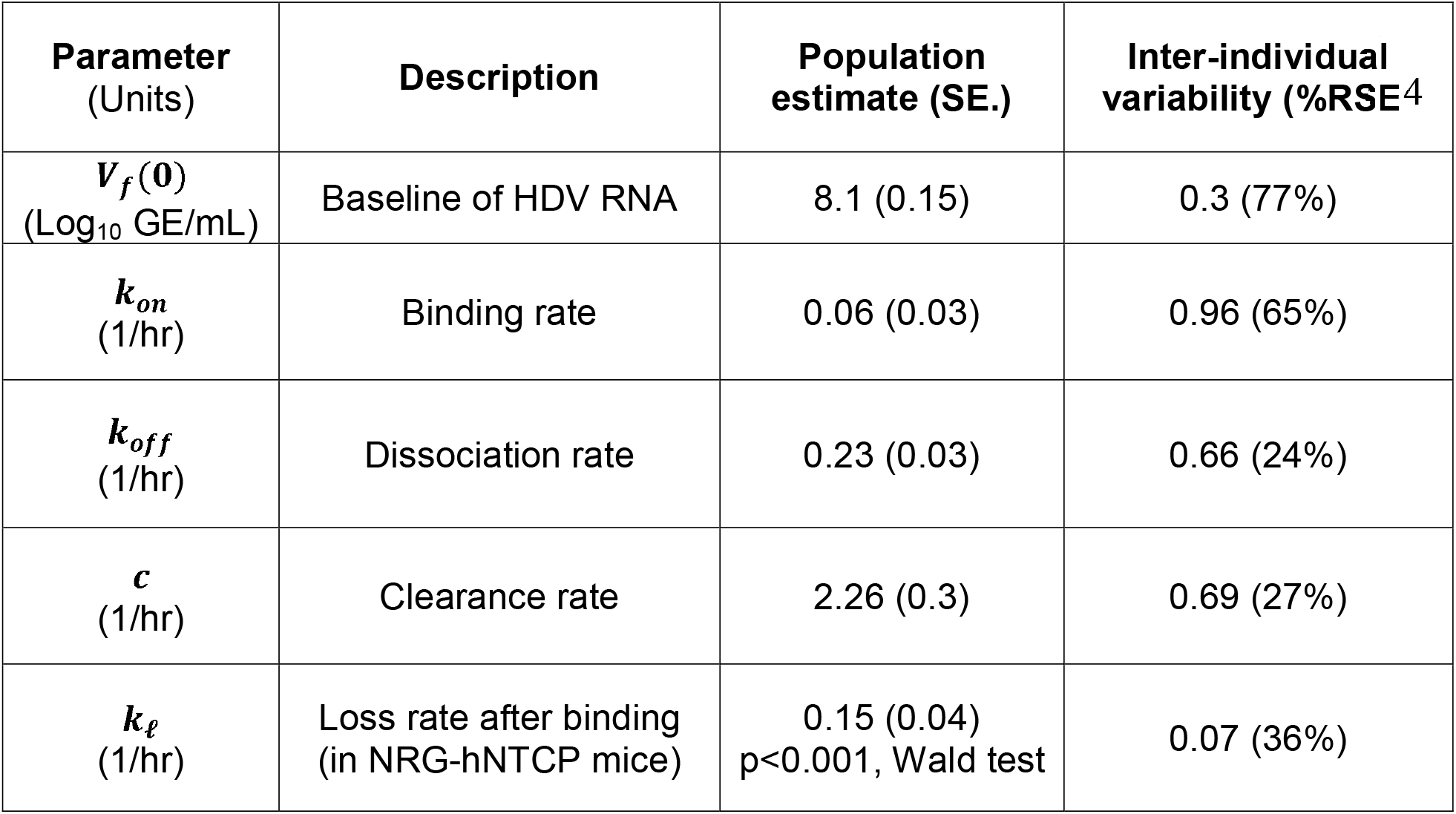
Mathematical model parameter estimates for the NRG and NRG-hNTCP mice population. As explained in the text, parameter value for *k*_ℓ_ only applies for NRG-hNTCP mice (i.e. *k*_ℓ_ *=0* for NRG mice). The table shows the MLE values for the fixed effects (population values) and the standard deviation of the random effects (inter-individual variation). SE: standard error; %RSE: % relative standard error.

**Figure 6.**
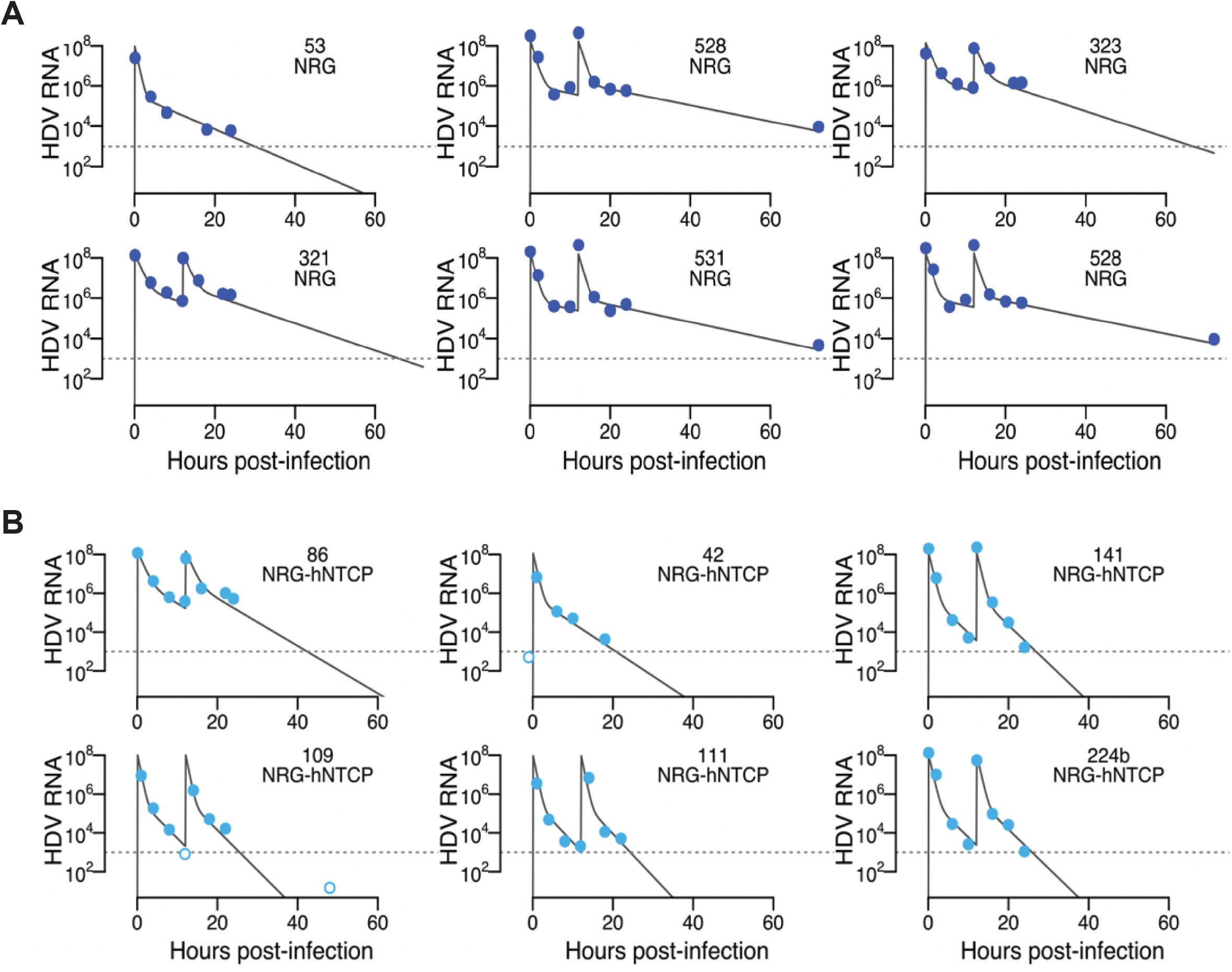
Best mathematical model fits of representative mice using the maximum likelihood estimates (MLE) of the population parameter distributions. Best model fits for some of the HDV concentrations (GE/mL) (**A**) from the NRG mice and (**B**) and NRG-hNTCP mice in the presence or absence of HDV re-inoculation at 12 hours after infection. Dotted horizontal line is the limit of quantification (LLoQ). Dark and light blue circles are data from NRG and NRG-hNTCP mice, respectively. Filled and empty circles are data over and below the LLoQ, respectively. Individual parameter estimates are in **Supp Table 3**. See methods for details of the assumptions for population distributions.

As presented in the previous section this model can recapitulate the biphasic decline in HDV concentration from all mice and can interpret the faster decline in the second phase for the NRG-hNTCP mice. From the best model fits we found that free HDV is cleared with rate 2.26 hour^-1^ (standard error, SE=0.32) equivalent to a half-life of 18 minutes (SE=2.4), binds to non-specific cells with rate 0.06 hour^-1^ (SE=0.03) and returned as free virus with rate 0.23 hour^-1^ (SE=0.03) (**Table 3**). The best model also shows that *k*_ℓ_ in the NRG-hNTCP mice is significantly greater than zero (p<0.001, Wald test) with rate 0.15 hour^-1^ (SE=0.04). This result implies that NRG-hNTCP mice have a loss of HDV bound to non-specific cells that does not become free virus.

## DISCUSSION

In this study, we utilized immunodeficient mice that we previously established (*23*) to characterize early HDV kinetics in immunocompetent versus immunodeficient backgrounds. These immunodeficient mice are non-obese diabetic recombinase activating gene 1 knockout (*Rag1*^*-/-*^) interleukin 2 receptor gamma chain deficient (*IL-2R*γ^*NULL*^) (NRG) mice, which lack functional natural killer cells, B and T lymphocytes (*32*), and express hNTCP (*23*). While hNTCP-transgenic mice are useful for studying HDV/HBV infection, we know little about early HDV kinetics in these mouse models. Therefore, we investigated the early kinetics of HDV infection in hNTCP transgenic mice on an immunocompetent (C57BL/6) or immunodeficient (NRG) background. We demonstrate that in all mice - irrespective of hNTCP expression and immune status - HDV RNA kinetics follows an unexcepted biphasic decline that is characterized by a sharp drop in the first phase and then a slower steady decrease until they reached LLoQ. Treating NRG-hNTCP mice with the HBV/HDV entry inhibitor, bulevirtide, suggests that viral entry is a minor contributor to HDV decline rates in NRG-hNTCP mice. Moreover, re-inoculating these mice with HDV 4 or 12 hpi still results in a biphasic decline of virus, suggesting that saturation of the interaction between HDV and its attachment factors – presumably HSPGs and/or hNTCP-is not achieved, which otherwise would have been expected to result in slowing viral clearance after the 2^nd^ inoculation. We employed roughly equal numbers of male (n=92, 48%) and female (n=100, 52%) mice and did not observe differences in HDV kinetics between genders of mice.

To increase the likelihood that the HDV inoculation resulted in high serum viremia detectable within the limits of our RT-qPCR assay and to ensure that the majority of target cells would be exposed to the virus, we began our experiments using a large dose of 10^9^ GE/mouse followed by 10^8^ GE/mouse in subsequent experiments, as our lab has previously done (*23*). Single inoculations of 10^9^ GE/mouse in immunocompetent (C57BL/6) and immunodeficient (NRG) mice either expressing hNTCP or not resulted in similar biphasic declines within the first 24 hours, with a sharp drop by 4 hpi followed by a slower decrease by 24 hpi. While the similar clearance rates were initially surprising, we reasoned that the adaptive immune response would not be primed within the first 24 hours of infection and therefore the initial HDV kinetics could resemble those in immunodeficient mice.

As the HDV RNA levels in NRG-hNTCP mice declined in a biphasic manner similarly to the other mouse cohorts, contrary to what was expected, we decided to analyze whether viral binding to NTCP was a factor to viral clearance in the blood through the use of bulevirtide, which binds to hNTCP and competitively inhibits HDV-receptor interactions. Since both hNTCP transgenic and non-transgenic C57BL/6 mouse cohorts exhibited viral decreases that were predicted due to their immunocompetency we did not include them in this experiment. Bulevirtide treatment on HDV-infected NRG-hNTCP mice did not affect the slopes of viral decline compared to those of HDV-infected NRG or NRG-hNTCP mice (**Fig 2B**), indicating that the hosts were able to eliminate the virus from circulation. This could be explained by bulevirtide binding non-specifically to HSPGs and/or binding to both endogenous mouse and human NTCP and thus resulting in similar kinetics of hNTCP and non-hNTCP mice.

To conjecture as to why the HDV RNA levels followed the same pattern in all three of these cohorts during single infection and BLV treatment, we explored the possibility that NTCP was saturated by viral binding. We thus injected mice at the end of the first and second phases, which corresponded with the timepoints of 4 and 12 hpi, respectively. Two main putative scenarios arise here: either viral RNA remains steady over time whereby the viral load will not decrease or there is a biphasic decline, in which there is no HDV-binding receptor saturation. Following 4 hpi re-inoculations, HDV RNA levels in NRG-hNTCP fell under the LLoQ by 8 hpi whereas RNA levels in NRG mice lasted over the LLoQ until 12 hpi. Moreover, HDV RNA levels in the 12 hpi re-inoculations of NRG-hNTCP mice fell under the LLoQ by 48 hpi whereas RNA levels of NRG mice hovered over the LLoQ by 72 hpi, indicating that re-inoculation of HDV in NRG-hNTCP mice consistently resulted in a faster second phase compared to NRG mice. This is notable because single inoculation of HDV in these mice yielded similar biphasic declines in NRG-hNTCP mice as in NRG, C57BL/6, and C57BL/6 mice. Moreover, re-inoculation of C57BL/6 and C57BL/6 -hNTCP mice emulated NRG kinetics, as the HDV RNA levels hovered above the LLoQ by 24 hpi (**Supp Fig 1**). HDV RNA levels in C57BL/6-hNTCP mice notably did not resemble those of NRG-hNTCP mice. We conjecture that this could possibly be due to clearance of the virus in the serum before the virus can attach to hNTCP. In contrast, in NRG-hNTCP mice, which are deficient in lytic complement and lack functional NK-, B and T cells, their immunocompromise status might delay viral clearance and thus allow binding of the virus to hNTCP. Thus, the steeper second phase viral decline found in NRG-hNTCP mice compared to NRG mice suggests a role of hNTCP in viral clearance from circulation.

We know that kinetics of the second phase decline are not influenced by the production and release of virions in the bloodstream, as the mice do not produce HBV or HBsAg, nor should there be a significant amount of HDV in the blood occurring from HDV-infected hepatocyte death (*23*). We therefore developed a mathematical model that assumes free-roaming HDV virions in the blood that can be bound to cells non-specifically without productive infection or viral replication, reminiscent of the binding compartment model used in (*33*). The model (**Fig. 5**) is able to recapitulate the biphasic decline of the observed HDV concentration and suggests that: the first phase of viral decline is explained by viral clearance rate from blood (c) and binding on-rate (k_on_) of the free-virus, and the second phase of decline by the dissociation of bound virus rate (k_off_) plus its loss rate (*k*_ℓ_) before dissociation only in hNTCP mice (either by disintegration or loss of bound virus that cannot return to circulation). Model fitting in NRG and NRG-hNTCP mice suggests that mice expressing the hNTCP receptor may sustain a significantly faster second phase of decline by virus loss after binding (i.e. *k*_ℓ_>0 in the model for the NRG-hNTCP mice). We speculate that this can be explained by two possible scenarios: the virus is entering hepatocytes or there may be yet unexplained ligand-virus interactions in transgenic hNTCP mice.

We estimated that the half-life of HDV in the bloodstream in our mouse model was 18 (SE 2.4) minutes. Interestingly, the half-life of HBV in chimeric urokinase-type plasminogen activator/severe combined immunodeficiency mice was found ∼3-fold longer i.e.,∼1 hour (*34*). HBV is a DNA virus and consequently is more stable in the bloodstream, whereas HDV, being an RNA virus, is less stable and could theoretically be cleared more rapidly by the host due to structural or nucleic acid degradation within the viral particles. Additionally, whether human hepatocytes clear HDV faster than mouse hepatocytes needs to be further investigated. The considerable difference in half-lives can also affect the process of antiviral administration as this clearance rate will need to be taken into account. Thus, delineating the mode and rate of viral decline will allow us to better understand natural HDV infection kinetics compared to anti-HDV therapeutic kinetics.

Elucidating HDV kinetics in these mouse models will further benefit from investigating several other aspects that were beyond the scope of this study. One such point is evaluating the effects of age on viral decline in these mouse models. Various studies have shown that young mice, those only 4 weeks of age or younger, are more susceptible to HDV infection and will develop chronic infections (*18, 35*). Additionally, including an HBsAg-producing mouse model will reveal whether HDV follows the same early kinetics or if newly synthesized virions could be found in the bloodstream 24 hpi. We did not inspect HDV replication in this study but this would help illustrate the ability for HDV to enter the hepatocytes within the first 24 hours of infection or if the virus requires more time. Future studies can also build upon the present study by evaluating bulevirtide treatment on HDV re-inoculations. This would provide insight into whether viral binding is the reason for decreased viral RNA in the bloodstream.

Altogether, this study demonstrates that adult C57BL/6, C57BL/6-hNTCP, NRG, and NRG-hNTCP mice undergo biphasic declines of viral RNA when inoculated with HDV in the absence of a helper virus, such as HBV. Remarkably, NRG-hNTCP mice displayed a more rapid second phase decline compared to NRG mice following double inoculation. Future studies, including characterization of the innate immune cell response and theoretical analysis will aid in the understanding of HDV-host dynamics and the effects of HDV antiviral therapy on early infection.

## Acknowledgments

We thank Stephan Urban (Heidelberg University, Germany) for providing the HDV-producing cell line, Huh7-END, and bulevirtide for our experiments and Preeti Dubey for initial mathematical modeling efforts.

## Abbreviations

NRG: (*NOD Rag1-/-IL2Rγ*^*NULL*^ *(NOD.Cg-Rag1*^*tm1Mom*^ *Il2rg*^*tm1Wjl/SzJ*^)
hNTCP: (human sodium taurocholate co-transporting polypeptide)
HDV: (hepatitis delta virus)
HBV: (hepatitis B virus)
tg: (transgenic)
LLOQ: (lower limit of quantification)
BAC: (bacterial artificial chromosome)
IACUC: (Institutional Animal Care and Use Committee)
GE: (genomic equivalents)
hpi: (hours post infection)
SEM: (standard error of the mean)
AIC: (Akaike Information Criteria)
ANCOVA: (Analysis of Covariance)
mpi: (minutes post infection)

