## supplementary information for "Hepatitis delta virus RNA decline post inoculation in human NTCP transgenic mice is biphasic"

for

**MATERIALS AND METHODS**

**Animals**

The NRG (*NOD Rag1-/- IL2Rγ^NULL^ (NOD.Cg-Rag1^tm1Mom^ Il2rg^tm1Wjl/SzJ^,* NRG)) and C57BL/6J mice were obtained from The Jackson Laboratory. hNTCP/BAC transgenic mice on the NRG and C57BL/6 background were generated as previously described (*23*) and can be obtained from The Jackson Laboratory (030533 and 030535 respectively, Bar Harbor, ME). Briefly, a bacterial artificial chromosome (BAC) expressing part of the human NTCP gene was microinjected into mice on the NRG and C57BL/6 background to create hNTCP/BAC transgenic mice. We employed both male (n=92, 48%) and female (n=100, 52%) mice in this study indiscriminately. Mice were housed in a cage system with a light/dark cycle. All in vivo experiments were conducted under a protocol (#3063) approved by the Institutional Animal Care and Use Committee (IACUC) of Princeton University. We utilized the ARRIVE1 reporting guidelines (*28*).

**Production of HDV stocks**

To generate HDV stocks for this study, Huh7-END cells (*29*) (kindly provided by Stephan Urban, University of Heidelberg) were grown and selected in DMEM supplemented with 5% (vol/vol) FBS, penicillin/streptomycin, G418 100 mg/mL, puromycin 10 mg/mL, and blasticidin 5 mg/mL. For HDV collection, the cells were grown in DMEM supplemented with 5% FBS, 1% penicillin/streptomycin, and 0.5% (vol/vol) DMSO. Supernatants were harvested every three days for 28 days and were sterile filtered through a 0.22 μm filter (Millipore, Burlington, MA) and concentrated 100-fold using a stir-cell concentrator (Millipore, Burlington, MA). The concentrated virus was then aliquoted into cryovial tubes and cryopreserved at -80 °C.

**Mouse injections**

Mouse injections with HDV were administered through intravenous injection into the tail vein with 1 x 10^9^ genomic equivalents (GE) or 1 x 10^8^ GE of HDV aliquoted in a volume of 200 μL per mouse. Mice doubly injected with HDV were injected at time zero and subsequently at 4 hours post infection (hpi) or 12 hpi. Bulevirtide was kindly gifted by Stephan Urban, University of Heidelberg. Bulevirtide was administered through subcutaneous injection at 2 mg/kg per mouse. Mice from all cohorts were bled submandibularly at the indicated time points and liver tissues were collected.

**RNA isolation from serum**

100 μL of blood was collected per mouse and was centrifuged at 3,500 rpm for 10 minutes at 4°C. Serum was extracted and aliquoted into microcentrifuge tubes and frozen at -80°C until further use. HDV RNA was isolated from 50 μL of serum utilizing the Zymo viral RNA isolation kit (Zymo, Irvine, CA) and was eluted in RNase-free and DNase-free water.

**Quantification of HDV RNA using RT-qPCR**

Quantification of HDV RNA was performed using a Quanta qScript one-step RT-qPCR kit (QuantaBio, Beverly, MA). 5 μL of RNA isolated from mouse serum or liver, HDV primers (forward: TGGACGTGCGTCCTCCT; reverse: TCTTCGGGTCGGCATGG; Eton Bioscience, San Diego, CA), HDV primer probe (ATGCCCAGGTCGGAC; Integrated DNA Technologies, San Jose, CA) and Quanta qScript qPCR master-mix (QuantaBio, Beverly, MA) were aliquoted into a 96-well plate for a total reaction volume of 20 μL. RNA samples were run in duplicate. RT-qPCR was done on a Step One Plus qPCR machine (LifeTechnologies, Carlsbad, CA) with the following program: 55°C for 20 minutes, 95°C for 3 minutes, 45 cycles (95°C for 20 seconds, 58°C for 45 seconds, and 72°C for 30 seconds).

**Kinetic analysis**

Kinetic analyses were conducted with Python 3.9 and statsmodels version 0.13.2. All values below the LLoQ were right-censored to LLoQ. For the purposes of understanding the kinetics dependent on each subset of mice used in this experiment, mice were agglomerated for kinetic analysis based on immunocompetence (NRG and C57BL/6), presence of the hNTCP receptor, and administration of bulevirtide. Further, kinetics for the first inoculation were assumed to be identical regardless of re-inoculation delay, so first-inoculation kinetics include all mice up until the point of their second inoculation (if at all). Distinctions between phases of decline were defined as a minimum 2-fold change in slope. Kinetics plots depict the median value at each time point, and the linear regressions were visualized with 95% confidence intervals (95% CI).

**Model fitting and parameter estimation**

We fit the ordinarily differential equation model in **Fig 5B** using a nonlinear mixed-effects approach. In this approach we assumed the log_10_ of the free HDV viral load in mouse $k$ time $j,$ is assumed to be $\log_{10} y_{j,k}=\log_{10} V_{f}(t_{j},x_{k})+\epsilon$, with $V_{f}$ the solution of the differential equation model (**Fig 5B**) for free virus at time $t_{j}$ with animal-specific parameter values $x_{k}$ and $\epsilon$ the error of the log_10_ of the HDV measurements with distribution $\mathcal{N}(0,\sigma_{v}^{2})$. The mixed-effects model assumes that for mouse $k$ the parameters $\kappa_{on_{k}},$ $k_{off_{k}}$ and $c_{k}$ come from distributions of the from $x_{k}=\hat{x}e^{\eta_{k}}$ with $\eta_{k}\mathcal{\sim N}(0,\omega_{x}^{2})$ being $\hat{x}$ fixed effects and $\omega_{x}$ the standard deviation of the random effects. We also assumed, for mouse $k$, that $V_{f_{k}}\left( t_{in} \right)={10}^{r+\eta_{r,k}}$with $\eta_{r}\mathcal{\sim N}(0,\omega_{r}^{2})$ and $t_{in}$ as the different times for HDV inoculation (0, 4 or 12 hours). Parameter $k_{i}$ was equal to zero for all of the non-transgenic mice and had population distribution $\log k_{i}\mathcal{\sim N(}\hat{k}_{i},\omega_{k_{i}}^{2})$ with $\hat{k}_{i}>0$for the transgenic hNTCP mice population.

We performed maximum likelihood estimation of the measurement error, fixed effects, standard deviation of the random effects and covariates for all different instances of the model using the Stochastic Approximation of the Expectation Maximization (SAEM) with the software Monolix. Monolix also computes %RSE is the relative standard error from the Fisher Information Matrix and quantifies the identifiability of a parameter with the available data—%RSE greater than 50% means that the parameter might not be identifiable. We assumed that $t_{j}=0$ represents the time of the first HDV infection.

**Statistical analysis**

All statistical analyses were conducted with GraphPad Prism software and in Python 3.9 with SciPy version 1.7.3. We tested whether covariates are significantly different than zero in the mathematical model fits using the Wald test in the Monolix software. Data are presented as means and the standard error of the mean (SEM). *P* values of ≤0.05 were noted as being statistically significant. To compare mathematical models, we used the Akaike Information Criteria (AIC) assuming that if the difference between the AICs of two models ΔAIC<4, the more complex model does not improve parsimony. In cases where multiple mice were agglomerated as a single time-series, statistical comparison of slopes was performed with the Analysis of Covariance (ANCOVA) test.

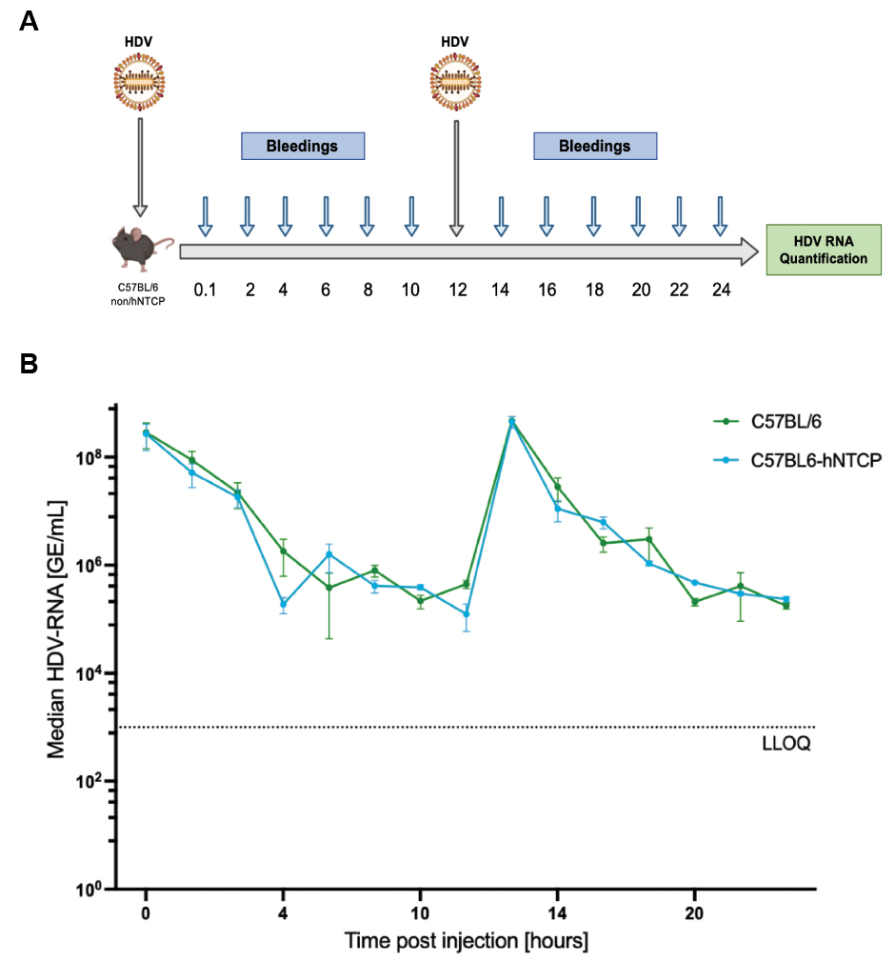

**Supplementary Figure 1. Re-infection of HDV in C57BL/6 and C57BL/6-hNTCP mice 12 hours post initial infection.**

(**A**) Schematic of C57BL/6 and NRG non-tg or tg mice infected with HDV and bled every two hours for the first 24 hours. Viral RNA was quantified from the serum by RT-qPCR. (**B**) Serum HDV RNA quantification over the first 24 hours of infection in C57BL/6 and C57BL/6-hNTCP mice. LLOQ indicates the lower limit of quantitation. Error bars are represented as standard error of the mean. Schematic created with Biorender.com.

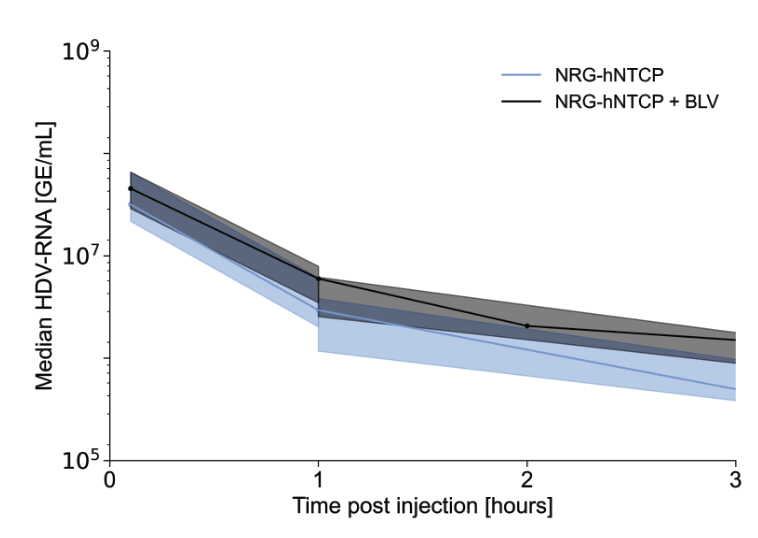

**Supplementary Figure 2. Slight clearance delay in NRG-hNTCP mice pre-treated with bulevirtide compared to untreated NRG-hNTCP mice.** HDV RNA levels in NRG-hNTCP mice declined somewhat more than NRG-hNTCP mice that were pre-treated with bulevirtide within the first few hours post inoculation of HDV. Shaded regions represent interquartile range.

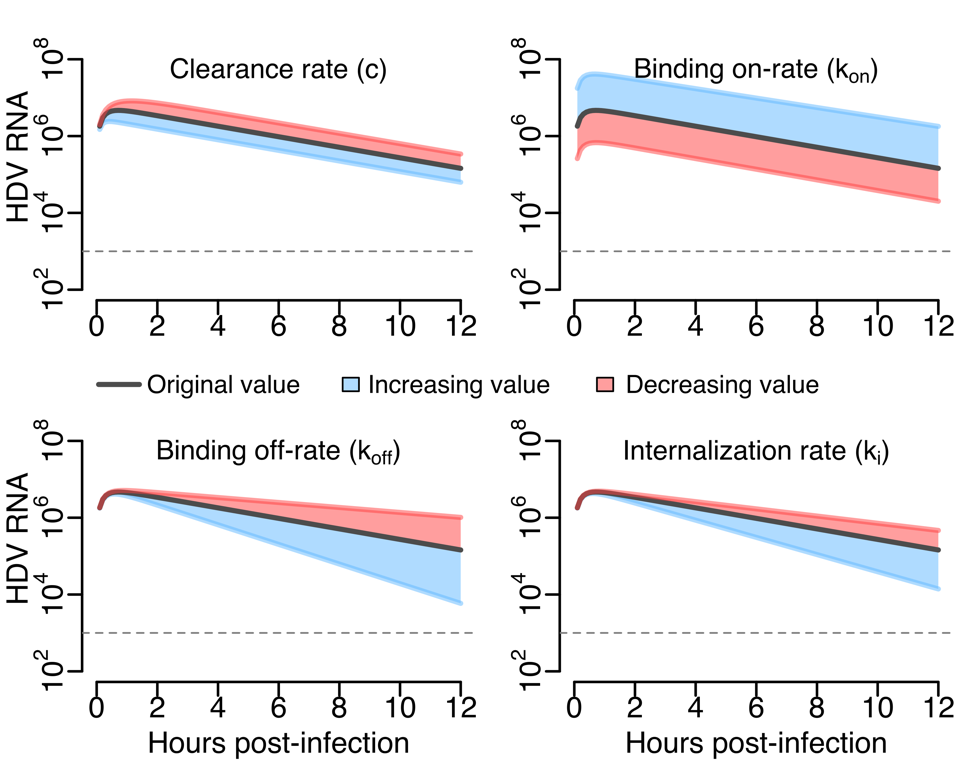

**Supplementary Figure 3. Mathematical model used to describe reinfection kinetics.**

Simulations of the model in **Fig 5A and 5B** for the cell-bound virus, V_b_. Model simulations indicate that the peak in cell-bound virus (V_b_) is inversely correlated with changes in clearance rate (top-left) and positively correlated with changes in binding on-rate (top-right). The second phase of decline is governed by the off-rate (right) and internalization constants (bottom-right). Model parameters in each graph are fixed at: $k_{on}$ = 0.07 hour^-1^, $k_{off}$= 0.22 hour^-1^, $k_{\mathcal{l}}$= 0.1 hour^-1^, $c$ = 3.71 hour^-1^ (black curve). Adjustments made to each parameter are shown in red (underestimate) and blue (overestimate), for illustrative purposes.

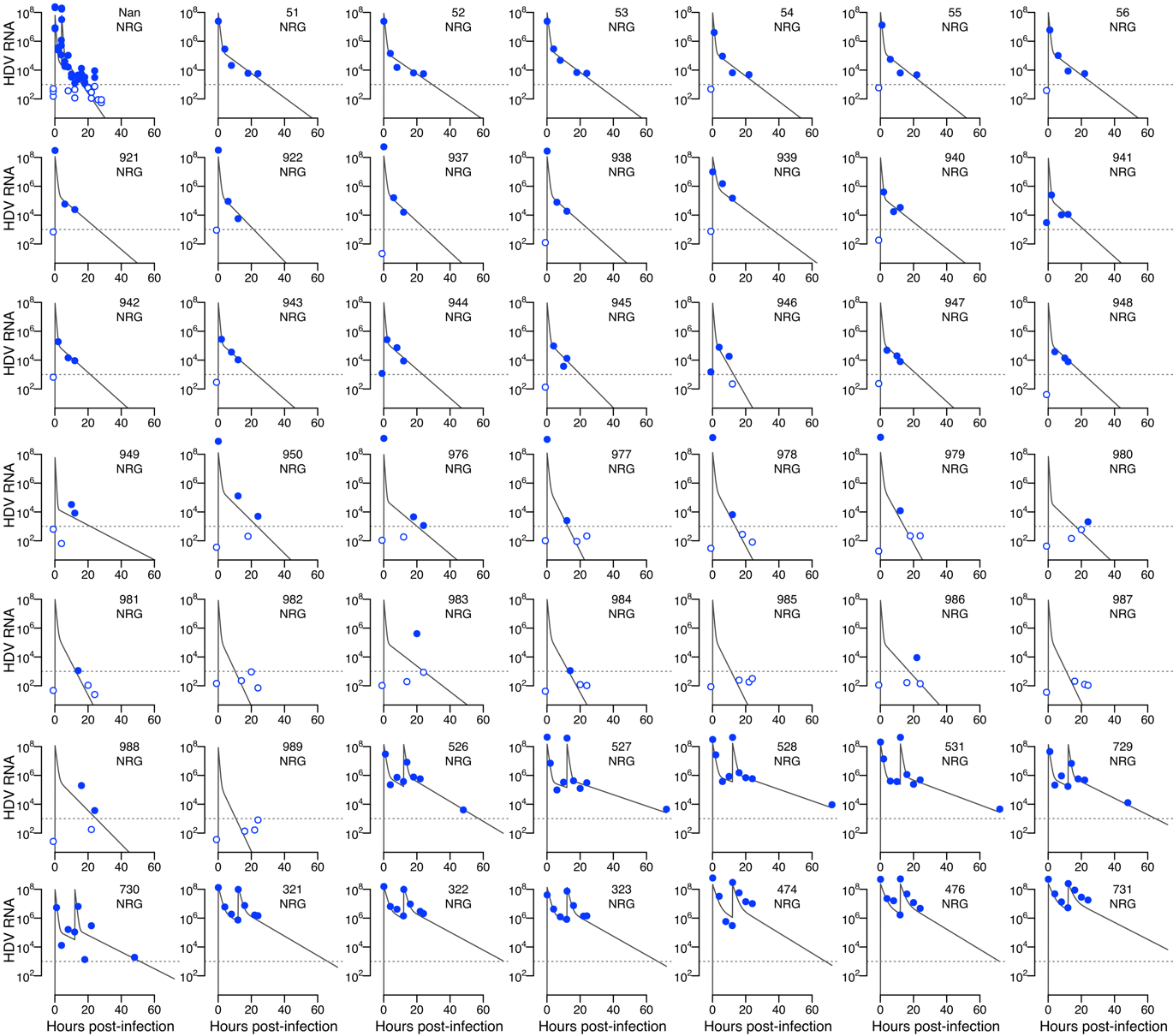

**Supplementary Figure 4. Mathematical model fits to the HDV viral loads from the NRG mice population.** Mathematical model fits of HDV concentrations (GE/mL) for individual mice in the NRG mouse cohort in the presence or absence of HDV re-inoculation. Dotted horizontal line is the limit of quantification (LLOQ). Filled and empty circles are data over and below the LLOQ, respectively.

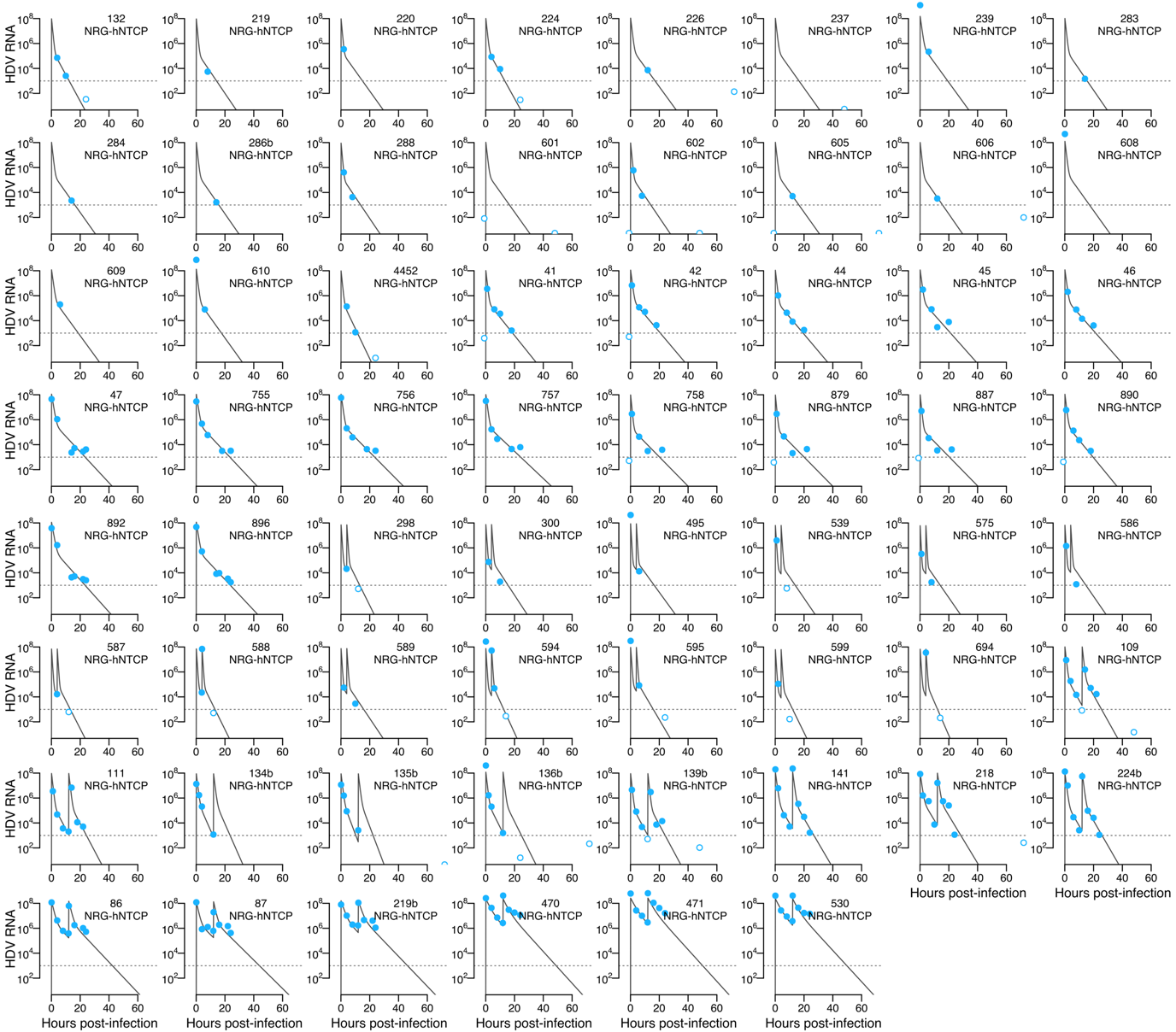

**Supplementary Figure 5. Mathematical model fits to the HDV viral loads from the NRG-hNTCP mice population.** Mathematical model fits of HDV concentrations (GE/mL) for individual mice in the NRG mouse cohort in the presence or absence of HDV re-inoculation. Dotted horizontal line is the limit of quantification (LLOQ). Filled and empty circles are data over and below the LLOQ, respectively.

| **ID** | $\mathbf{lo}\mathbf{g}_{\mathbf{10}}\boldsymbol{V}_{\boldsymbol{f}}\boldsymbol{(0)}$ | $\boldsymbol{k}_{\boldsymbol{on}}$ | $\boldsymbol{k}_{\boldsymbol{off}}$ | $\boldsymbol{c}$ |
| --- | --- | --- | --- | --- |
| **Unidentified** | 7.97 | 0.04 | 0.37 | 3.85 |
| **51** | 8.06 | 0.06 | 0.21 | 2.27 |
| **52** | 8.05 | 0.05 | 0.19 | 2.50 |
| **53** | 8.07 | 0.06 | 0.21 | 2.15 |
| **54** | 8.09 | 0.06 | 0.21 | 2.80 |
| **55** | 8.10 | 0.05 | 0.21 | 2.51 |
| **56** | 8.10 | 0.06 | 0.21 | 2.61 |
| **921** | 8.16 | 0.05 | 0.23 | 2.43 |
| **922** | 8.15 | 0.05 | 0.27 | 2.56 |
| **937** | 8.19 | 0.06 | 0.25 | 2.27 |
| **938** | 8.16 | 0.05 | 0.24 | 2.41 |
| **939** | 8.09 | 0.08 | 0.22 | 1.48 |
| **940** | 8.10 | 0.06 | 0.22 | 2.82 |
| **941** | 8.08 | 0.05 | 0.24 | 3.11 |
| **942** | 8.08 | 0.05 | 0.24 | 3.21 |
| **943** | 8.10 | 0.06 | 0.23 | 2.97 |
| **944** | 8.11 | 0.06 | 0.23 | 2.95 |
| **945** | 8.08 | 0.05 | 0.26 | 2.81 |
| **946** | 8.08 | 0.05 | 0.45 | 3.00 |
| **947** | 8.09 | 0.05 | 0.24 | 2.81 |
| **948** | 8.08 | 0.05 | 0.24 | 2.92 |
| **949** | 7.97 | 0.03 | 0.15 | 4.72 |
| **950** | 8.20 | 0.05 | 0.26 | 2.36 |
| **976** | 8.14 | 0.03 | 0.21 | 3.62 |
| **977** | 8.20 | 0.05 | 0.52 | 2.70 |
| **978** | 8.22 | 0.05 | 0.50 | 2.49 |
| **979** | 8.23 | 0.05 | 0.48 | 2.42 |
| **980** | 8.01 | 0.04 | 0.24 | 3.94 |
| **981** | 8.10 | 0.05 | 0.51 | 2.66 |
| **982** | 8.06 | 0.04 | 0.55 | 3.32 |
| **983** | 8.07 | 0.05 | 0.21 | 2.97 |
| **984** | 8.10 | 0.05 | 0.47 | 2.74 |
| **985** | 8.07 | 0.04 | 0.52 | 3.13 |
| **986** | 8.06 | 0.04 | 0.28 | 3.18 |
| **987** | 8.07 | 0.04 | 0.54 | 3.13 |
| **988** | 8.15 | 0.06 | 0.27 | 1.98 |
| **989** | 8.07 | 0.04 | 0.55 | 3.16 |
| **526** | 8.21 | 0.09 | 0.19 | 1.62 |
| **527** | 8.25 | 0.06 | 0.14 | 1.85 |
| **528** | 8.29 | 0.07 | 0.15 | 1.38 |
| **531** | 8.25 | 0.07 | 0.15 | 1.53 |
| **729** | 8.21 | 0.08 | 0.17 | 1.67 |
| **730** | 8.08 | 0.04 | 0.16 | 2.30 |
| **321** | 8.24 | 0.09 | 0.20 | 0.97 |
| **322** | 8.26 | 0.10 | 0.20 | 0.90 |
| **323** | 8.18 | 0.09 | 0.20 | 0.99 |
| **474** | 8.37 | 0.07 | 0.20 | 0.76 |
| **476** | 8.41 | 0.09 | 0.22 | 0.67 |
| **731** | 8.43 | 0.10 | 0.21 | 0.56 |

**Supplementary Table 1.** Mathematical model, individual parameter estimates for the NRG mice population. As specified in the text, the value of $k_{\mathcal{l}}$ for this group of mice was zero.

| **ID** | $\mathbf{lo}\mathbf{g}_{\mathbf{10}}\boldsymbol{V}_{\boldsymbol{f}}\boldsymbol{(0)}$ | $\boldsymbol{k}_{\boldsymbol{on}}$ | $\boldsymbol{k}_{\boldsymbol{off}}$ | $\boldsymbol{c}$ | $\boldsymbol{k}_{\mathcal{l}}$ |
| --- | --- | --- | --- | --- | --- |
| **132** | 8.10 | 0.05 | 0.33 | 2.67 | 0.17 |
| **219** | 8.09 | 0.05 | 0.24 | 2.74 | 0.16 |
| **220** | 8.11 | 0.06 | 0.23 | 2.84 | 0.15 |
| **224** | 8.12 | 0.05 | 0.33 | 2.42 | 0.16 |
| **226** | 8.13 | 0.06 | 0.22 | 2.18 | 0.15 |
| **237** | 8.13 | 0.06 | 0.23 | 2.27 | 0.15 |
| **239** | 8.26 | 0.06 | 0.23 | 1.84 | 0.15 |
| **283** | 8.12 | 0.06 | 0.24 | 2.40 | 0.15 |
| **284** | 8.12 | 0.06 | 0.23 | 2.30 | 0.15 |
| **286b** | 8.12 | 0.06 | 0.24 | 2.37 | 0.15 |
| **288** | 8.08 | 0.05 | 0.24 | 2.90 | 0.16 |
| **601** | 8.13 | 0.06 | 0.23 | 2.27 | 0.15 |
| **602** | 8.09 | 0.05 | 0.24 | 2.74 | 0.16 |
| **605** | 8.13 | 0.06 | 0.23 | 2.28 | 0.15 |
| **606** | 8.12 | 0.06 | 0.24 | 2.40 | 0.15 |
| **608** | 8.20 | 0.06 | 0.23 | 2.19 | 0.15 |
| **609** | 8.16 | 0.07 | 0.23 | 1.86 | 0.15 |
| **610** | 8.22 | 0.06 | 0.23 | 2.13 | 0.15 |
| **4452** | 8.10 | 0.05 | 0.40 | 2.56 | 0.17 |
| **41** | 8.12 | 0.08 | 0.20 | 2.31 | 0.13 |
| **42** | 8.14 | 0.08 | 0.19 | 1.99 | 0.13 |
| **44** | 8.14 | 0.07 | 0.19 | 2.25 | 0.13 |
| **45** | 8.15 | 0.07 | 0.18 | 1.93 | 0.13 |
| **46** | 8.16 | 0.07 | 0.18 | 1.95 | 0.13 |
| **47** | 8.12 | 0.06 | 0.17 | 1.54 | 0.12 |
| **755** | 8.11 | 0.07 | 0.17 | 1.66 | 0.12 |
| **756** | 8.13 | 0.07 | 0.16 | 1.88 | 0.11 |
| **757** | 8.10 | 0.07 | 0.15 | 1.92 | 0.11 |
| **758** | 8.11 | 0.07 | 0.16 | 2.63 | 0.12 |
| **879** | 8.10 | 0.07 | 0.16 | 2.67 | 0.12 |
| **887** | 8.11 | 0.07 | 0.16 | 2.48 | 0.12 |
| **890** | 8.13 | 0.07 | 0.20 | 2.11 | 0.13 |
| **892** | 8.12 | 0.06 | 0.18 | 1.43 | 0.13 |
| **896** | 8.13 | 0.07 | 0.17 | 1.63 | 0.12 |
| **298** | 8.02 | 0.04 | 0.37 | 3.99 | 0.17 |
| **300** | 8.04 | 0.04 | 0.25 | 3.82 | 0.16 |
| **495** | 8.13 | 0.05 | 0.22 | 4.43 | 0.15 |
| **539** | 7.98 | 0.03 | 0.22 | 4.05 | 0.17 |
| **575** | 8.01 | 0.04 | 0.23 | 5.13 | 0.16 |
| **586** | 8.00 | 0.03 | 0.23 | 4.25 | 0.16 |
| **587** | 8.02 | 0.04 | 0.35 | 4.06 | 0.17 |
| **588** | 8.02 | 0.04 | 0.37 | 3.97 | 0.17 |
| **589** | 8.04 | 0.04 | 0.24 | 3.92 | 0.16 |
| **594** | 8.06 | 0.04 | 0.41 | 4.24 | 0.17 |
| **595** | 8.13 | 0.05 | 0.29 | 3.65 | 0.16 |
| **599** | 7.97 | 0.03 | 0.37 | 4.04 | 0.18 |
| **694** | 8.00 | 0.04 | 0.45 | 4.00 | 0.17 |
| **109** | 8.11 | 0.05 | 0.32 | 2.33 | 0.16 |
| **111** | 8.07 | 0.04 | 0.34 | 2.30 | 0.17 |
| **134b** | 8.04 | 0.06 | 0.45 | 2.15 | 0.17 |
| **135b** | 8.04 | 0.06 | 0.56 | 2.21 | 0.17 |
| **136b** | 8.18 | 0.05 | 0.38 | 2.28 | 0.17 |
| **139b** | 8.06 | 0.04 | 0.33 | 2.45 | 0.17 |
| **141** | 8.18 | 0.05 | 0.30 | 1.95 | 0.16 |
| **218** | 8.07 | 0.08 | 0.30 | 1.77 | 0.16 |
| **224b** | 8.10 | 0.04 | 0.30 | 2.02 | 0.17 |
| **86** | 8.21 | 0.11 | 0.19 | 0.99 | 0.11 |
| **87** | 8.18 | 0.13 | 0.18 | 1.06 | 0.10 |
| **219b** | 8.24 | 0.11 | 0.20 | 0.71 | 0.11 |
| **470** | 8.37 | 0.08 | 0.23 | 0.48 | 0.13 |
| **471** | 8.44 | 0.07 | 0.23 | 0.44 | 0.13 |
| **530** | 8.39 | 0.08 | 0.23 | 0.47 | 0.13 |

**Supplementary Table 2.** Mathematical model, individual parameter estimates for the NRG-hNTCP mice population.
